# Nonlinear Functional Network Connectivity in Resting Fmri Data

**DOI:** 10.1101/2021.07.20.452982

**Authors:** S. M. Motlaghian, A. Belger, J. R. Bustillo, J. M. Ford, K. Lim, D. H. Mathalon, B. A. Mueller, D. O’Leary, G. Pearlson, S. G. Potkin, A. Preda, T.G. van Erp, V. D. Calhoun

**Affiliations:** Tri-institutional Center for Translational Research in Neuroimaging and Data Science (TReNDS), Georgia State, Georgia Tech, Emory, Atlanta, Georgia, USA; Department of Psychiatry, University of North Carolina, Chapel Hill, NC, USA; Department of Psychiatry, University of New Mexico Albuquerque, NM, USA; Department of Psychiatry, University of California San Francisco, San Francisco, CA, USA; San Francisco VA Medical Center, San Francisco, CA, USA; Department of Psychiatry, University of Minnesota, Minneapolis, MN, USA; Department of Psychiatry, University of Iowa, Iowa City, IA, USA; Department of Psychiatry and Neurobiology, Yale School of Medicine, New Haven, CT, USA; Department of Psychiatry and Human Behavior, University of California Irvine, Irvine, CA, USA

**Keywords:** Mutual information, functional network connectivity, time courses, nonlinear functional network connectivity

## Abstract

In this work, we focus on explicitly nonlinear relationships in functional networks. We introduce a technique using normalized mutual information (MI), that calculates the nonlinear correlation between different brain regions. We demonstrate our proposed approach using simulated data, then apply it to a dataset previously studied in (Damaraju et al., 2014). This resting-state fMRI data included 151 schizophrenia patients and 163 age- and gender-matched healthy controls. We first decomposed these data using group independent component analysis (ICA) and yielded 47 functionally relevant intrinsic connectivity networks. Our analysis showed a modularized nonlinear relationship among brain functional networks that was particularly noticeable in the sensory and visual cortex. Interestingly, the modularity appears both meaningful and distinct from that revealed by the linear approach. Group analysis identified significant differences in nonlinear dependencies between schizophrenia patients and healthy controls particularly in visual cortex, with controls showing more nonlinearity in most cases. Certain domains, including cognitive control, and default mode, appeared much less nonlinear, whereas links between the visual and other domains showed evidence of substantial nonlinear and modular properties. Overall, these results suggest that quantifying nonlinear dependencies of functional connectivity may provide a complementary and potentially important tool for studying brain function by exposing relevant variation that is typically ignored.

Further, we propose a method that captures both linear and nonlinear effects in a ‘boosted’ approach. This method increases the sensitivity to group differences in comparison to the standard linear approach, at the cost of being unable to separate linear and nonlinear effects.

## Introduction

Functional connectivity (FC) has been widely used to assess linear dependencies among brain activity (Bastos & Schoffelen, 2016; Friston, 2011; Sala-Llonch, Bartrés-Faz, & Junqué, 2015; van den Heuvel & Hulshoff Pol, 2010). Allen et al. (Allen et al., 2011) applied an approach that characterized whole-brain functional network connectivity (FNC, i.e., FC between coherent intrinsic networks) to a large dataset. Most functional connectivity studies concentrate on linear relationships that have many benefits but also some limitations such as ignoring nonlinear contributions. One of the most widely used methods for assessing functional connectivity is the correlation coefficient, which is easy to calculate and interpret for both positive and negative correlation. However, the correlation coefficient measures only linear dependency. There has been little work studying explicitly nonlinear relationships in functional connectivity.

In the current study, we were interested in evaluating the degree to which nonlinear relationships exist among brain regions in a functional connectivity context. To do this we focused on the use of mutual information (MI), an information theoretic approach that has the advantage of being capable of measuring both linear and nonlinear dependencies. Early work evaluated MI as a way to capture more general relationships (V. Calhoun, Kim, & Pearlson, 2003). More recently, alternative metrics for functional connectivity, including MI, have been explored (Mohanty, Sethares, Nair, & Prabhakaran, 2020; Sundaram et al., 2020; Tedeschi et al., 2005; Tsai et al., 1999; Wang et al., 2015; Zhang, Muravina, Azencott, Chu, & Paldino, 2018). However, to our knowledge, we are the first group to assess the explicitly nonlinear relationships among brain networks to evaluate their unique aspects relative to the linear relationships.

We have developed an approach that explores the nonlinear relationships in functional connectivity after first removing the linear relationships via regression. This is followed by an estimation of the mutual information among the residual time courses. In order to assess whether the nonlinear relationships were potentially meaningful we first focus on whether the resulting FNC matrices exhibit modular relationships, consistent with functional integration. Secondly, we evaluated whether the nonlinear FNC show group differences in a dataset consisting of resting fMRI data collected from schizophrenia patients and healthy controls (Damaraju et al., 2014). To do this, the MI among the functional brain networks in patients and controls were assessed and compared after removal of the linear dependencies.

## Materials and Methods

### 2.1 Participants and Preprocessing

In this work, we use the fBIRN dataset which has been analyzed previously in (Damaraju et al.). The final curated dataset consisted of 163 healthy participants (mean age 36.9, 117 males; 46 females) and 151 age- and gender-matched patients with schizophrenia (mean age 37.8; 114 males, 37 females). Eyes-closed resting state fMRI data were collected at 7 sites across the United State (Keator et al., 2016). Informed consent was obtained from all subjects prior to scanning in accordance with the Internal Review Boards of corresponding institutions. With a TR of 2s on 3T scanners, 162 volumes of echo-planar imaging blood oxygenation level-dependent (BOLD) fMRI data were captured. Imaging data of one site was captured on General Electric Discovery MR750 scanner and the rest of the six sites were collected on Siemens Tim Trio System. Resting-state fMRI scans were acquired using a standard gradient-echo echo-planar imaging paradigm: FOV of 220 × 220 mm (64 × 64 matrices), TR = 2 s, TE = 30 ms, FA = 770, 162 volumes, 32 sequential ascending axial slices of 4 mm thickness and 1 mm skip.

Data preprocessed by using several toolboxes such as AFNI, SPM, GIFT. Rigid body motion correction using the INRIAlign (Freire & Mangin, 2001) toolbox in SPM to correct head motion was applied. To remove the outliers, the AFNI3s 3dDespike algorithm was performed. Then fMRI data were resampled to 3 mm^3^ isotropic voxels. Then data were smoothed to 6 mm full width at half maximum (FWHM) using AFNI3s BlurToFWHM algorithm and each voxel time course was variance normalized. During the curation process, subjects with larger movement were excluded from the analysis to mitigate motion effects. For more details please see (Damaraju et al.).

### 2.2. Postprocessing

The GIFT (http://trendscenter.org/software/gift) implementation of Group-level Spatial ICA was used to estimate 100 functional networks as ICA components. A subject-specific data reduction step was first used to reduce 162 time point data into 100 directions of maximal variability using principal component analysis. Next, the infomax approach (Bell & Sejnowski, 1995) was used to estimate one hundred maximally independent components from the group PCA reduced matrix. For stability of estimation, the ICA algorithm was repeated multiple times and the most central run was selected as representative (Du, Ma, Fu, Calhoun, & Adalı, 2014). Finally aggregated spatial maps were estimated as the modes of component clusters. Subject specific spatial maps (SMs) and time courses (TCs) were obtained using the spatiotemporal regression back reconstruction approach (V. D. Calhoun, Adali, Pearlson, & Pekar, 2001; Erhardt et al., 2011) implemented in the GIFT software.

To label the components, regions of peak activation clusters for each specific spatial map were obtained. After ICA processing, to acquire regions of peak activation clusters, one sample t-test maps are taken for each SM across all subjects and then thresholded; also mean power spectra of the corresponding TCs were computed. The set of components as intrinsic connectivity networks (ICNs) is identified if their peak activation clusters fell within gray matter and showed less overlap with known vascular, susceptibility, ventricular, and edge regions corresponding to head motion. This resulted in 47 ICNs out of the 100 independent components. Running over 20 times ICASSO, the cluster stability/quality index for all except one ICNs was very high. After TCs were detrended and orthogonalized by considering estimated subject motion parameters, spikes were detected by AFNI3s 3dDespike algorithm and replaced by values of third order spline fit. For more detail see (Allen et al., 2012; Damaraju et al., 2014). The fBIRN dataset obtained after processing resulted in a matrix of 159 time points × 47 components × 314 subjects including 163 Control and 151 SZ subjects.

### 2.3. A Mutual Information Approach

While linear correlation is the most widely used measure to describe dependence, it can underestimate or completely miss nonlinear dependencies. An example to illustrate this shortfall is Anscombe’s Quartet (Anscombe, 1973) where four plots of various, non-random data points (linear dependence, quadratic dependence, straight line with outliers, and vertical line with outliers) are shown that have wildly different structure of dependence but the same correlation coefficient. To measure the explicitly nonlinear correlation between two TCs, the approach applied in this research is to remove the linear correlation and calculate the residual dependence.

The Pearson product moment correlation coefficient, ρ, of time courses *x* and *y* is

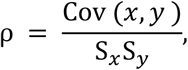

where *S*_*x*_ and *S*_*y*_ are respectively the sample standard deviations of *x* and *y*, and Cov (*x*, *y*) is the covariance between *x* and *y*. The correlation coefficient mainly measures the linear dependence between two distributions. However nonlinear dependence is not displayed in the value of the correlation coefficient. Recent statistical approaches have been proposed to measure the correlation without underestimating the nonlinear dependency. One of these methods, normalized mutual information (MI), measures both linear and nonlinear dependencies. The value of MI is determined by the formula

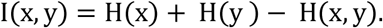

Where H(x) and, H(y) are marginal entropies and H(x, y) is the joint entropy.

The goal is to calculate only the nonlinear component of dependence. The approach we use for this is to measure the mutual information of the data after removal of the linear component of dependence. For given time courses *x* and *y*, fitting a linear model 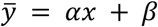 gives the linear correlation between *x* and *y*. Next, we cancel the linear effect by calculating 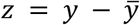. The nonlinear dependency of *x* and *z* is the same as *x* and *y*. Next, we can use I(*x*, *z*) to evaluate the nonlinear dependency of *x* and *y*.

### 2.4. Simulated Experiment

We applied the proposed method to simulated data to illustrate their use. In this experiment, we started with a vector say *x* of size 1000 × 1 where its components are from a random uniform distribution on [0 1]. Next, we formed three vectors *y*_1_, *y*_2_, and *y*_3_, such that each one has a particular relationship with *x*. Three different types of relationships are as follows: **Case I,** vector *y*_1_ has a purely linear relationship with *x*. **Case II,** we defined *y*_2_ to have a quadratic relationship and no linear correlation with *x*. That is *x* and *y*_2_ have only a nonlinear correlation. **Case III,** vector *y*_3_ has a combination of linear and nonlinear correlation with *x*. We also added zero-mean Gaussian noise to *y*_1_, *y*_2_, and *y*_3_ (**Fig. 1**).

**Fig. 1:**
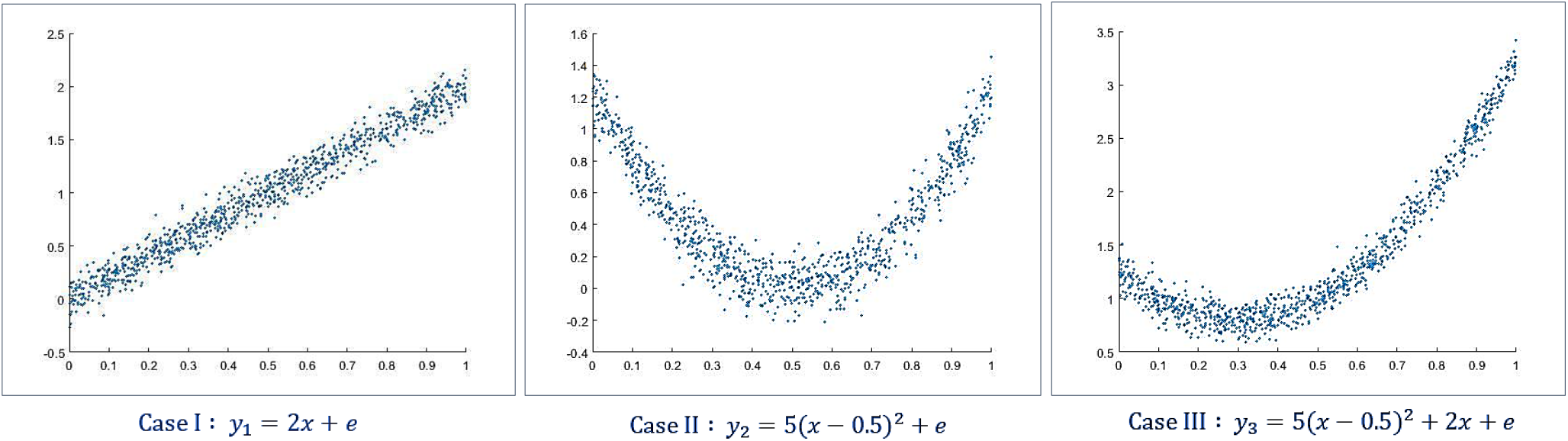
Three simulation cases for linear and nonlinear correlation between two vectors. Vector *x* has its components randomly derived from a uniform distribution on [0 1]. From left to right we have Case I, Case II and Case III such that in Case I, *y*_1_ = 2*x* + *e* (linear relationship between *x* and *y*_1_). In Case II, we have *y*_2_ = 5(*x* − 0.5)^2^ + *e* (nonlinear relationship between *x* and *y*_2_) and for Case III, *y*_3_ = 5(*x* − 0.5)^2^ + 2*x* + *e* (combination of linear and nonlinear relationships between *x* and *y*_3_). Noise *e* is a Gaussian distribution with a mean of zero.

We measured the relationship of (*x*, *y*_1_), (*x*, *y*_2_) and (*x*, *y*_3_) using both Pearson correlation and mutual information approaches. Pearson correlation takes value from −1 to 1. Briefly, −1 refers to a perfectly linear negative correlation and 1 shows a perfectly linear positive correlation. The mutual information we use in this work is normalized, taking a range of [0,1]. If the MI=0 this indicates no dependency and as two distributions increase their dependency, the mutual information value increases to a maximum of 1. Prior to computing correlation and MI, we implemented the procedure explained earlier to remove the linear correlation from *y*_1_, *y*_2_, and *y*_3_. Next, we calculated the Pearson correlation and mutual information for each pair as it shown in **Table 1**.

**Table 1.** Simulation of three cases, including Case I: linear correlation, Case II: Nonlinear correlation and, Case III: Linear and nonlinear correlation between two vectors. The contribution of two vectors in each case measured by Pearson correlation (Corr) and Mutual Information (MI). In this table Corr1 and MI1 show the correlation between the original data and Corr2 and MI2 show the correlation after removing the linear relationship. Range of Pearson correlation is [−1, 1] and mutual information is normalized [0,1]. As expected, the correlation is effectively zero after removal of linear effects. Results show that correlation completely misses the residual nonlinear dependencies, and that the MI approach is able to effectively capture the nonlinear relationships when they exist.

| | $Corr_1(x, \dots)$ | $Corr_2(x, \dots)$ | $MI_1(x, \dots)$ | $MI_2(x, \dots)$ |
| --- | --- | --- | --- | --- |
| Case I : $y_1 = 2x + e$ | 0.9847 | $1.64 \times 10^{-15}$ (i.e., 0) | 0.3809 | 0.0161 |
| Case II : $y_2 = 5(x - 0.5)^2 + e$ | -0.0271 | $-3.91 \times 10^{-17}$ (i.e., 0) | 0.2585 | 0.2582 |
| Case III : $y_3 = 5(x - 0.5)^2 + 2x + e$ | 0.8276 | $-1.61 \times 10^{-15}$ (i.e., 0) | 0.3139 | 0.2630 |

### 2.5. Quantifying Nonlinear Connectivity in fMRI Data

For each subject there are 47 ICA time courses of length 159. For each pair of time courses *x* and *y* we compute the traditional FNC, i.e., the linear correlation between all pairs *x* and *y*. We then compute the study mean FNC matrix by averaging over all 314 subjects included in this analysis.

We then fit a linear model to estimate the linear correlation between *x* and *y*. After this we remove the linear effect to study the remaining dependencies by updating *y* as: 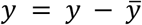. Finally, we calculate the nonlinear residual dependencies among functional network components in fMRI data via MI. This produces a matrix of 47 by 47 for each subject in which, the value in (*x*, *y*) entry shows the nonlinear dependencies calculated by MI method for *x* and *y*. Then the average over all 314 subjects is calculated. In order to examine whether these differences are significant, we compute a one-sample T-test comparing each MI value against the cell with the minimum MI average value in the matrix.

Next, we compare the dependencies between the schizophrenia patients and controls. Within each group, the linear effect is canceled, and the average MI is calculated over all subjects. We first evaluate whether there is modular structure in the FNC matrix. Using T-test for two samples, the differences of nonlinear correlation are calculated. Also, for false discovery rate (FDR) correction, all p-values were adjusted by the Benjamini-Hochberg correction method and threshold at a corrected p<0.05.

### 2.6. Boosted Approach

While we emphasize the unique information contained in the nonlinearities, future studies may wish to leverage both linear and nonlinear information together. To do this, we propose a method that include the information from both linear and nonlinear dependencies. This *boosted* approach is a combination of Pearson correlation and modified mutual information for quantifying nonlinear dependencies as described in *2.3. A Mutual Information Approach.* We define the boosted method as

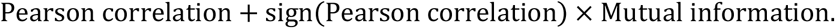

With this technique, we use the nonlinear information to boost the linear effects in the direction of the Pearson correlation. Thus, both linear and nonlinear correlation is considered and, the direction (which is not well defined in the nonlinear case) of the linear effect is protected. The approach we propose allows for multiple uses. One can focus on the nonlinear effects only, which as we show in this paper, may be interesting in and of themselves. Secondly, one can focus on capturing both linear and nonlinear effects in a ‘boosted’ approach. This appears to increase sensitivity to group differences beyond the standard linear approach, though it does not allow for separately of linear and nonlinear effects.

We assessed the linear correlation (Pearson correlation), nonlinear dependencies (modified mutual information *2.3. A Mutual Information Approach*) and both linear and nonlinear dependencies (Boosted) in schizophrenia patients and healthy controls components. Then separately for each method, T-test for two samples was applied and p-values were adjusted by the Benjamini-Hochberg correction method and threshold at a corrected p<0.05.

### 2.7. Joint Distributions

To further visualize the identified nonlinear relationships, we selected the five component pairs that have the most significant p-values in the T-test for group differences in the nonlinear dependence for HC-SZ. Then we constructed the difference in the joint distributions for each pair of time courses, comparing patients and controls.

## Results

### 3.1. Simulated Experiment

Three types of correlations: linear, nonlinear and combination of linear and nonlinear, are examined. The Pearson correlation and mutual information before and after removing linear dependency for each case are measured and reported in the **Table 1**.

The range of Pearson correlation is −1 to +1 and the range of mutual information for independent distributions is 0, and perfect dependency is 1. In Case I, where the two distributions have only a linear correlation, the Pearson correlation before removing the linear effect is close to one and after removing, both Pearson and mutual information are close to zero. In Case II, where the two distributions have a quadratic relationship, the Pearson correlation shows a low but non-zero correlation while the mutual information calculation shows a considerable correlation between the two distributions. After removing the linear effects, Person correlation is effectively zero while the mutual information is approximately the same before and after removing the linear effect. In Case III, where there are both linear and nonlinear correlation between two distributions, the Pearson correlation is significant before removing the linear effect and vanishes after canceling the linear correlation, while the mutual information only slightly decreased after removing linear correlation. This provides a straightforward demonstration of the fact that Pearson correlation is not able to capture purely nonlinear dependencies, while mutual information considers both linear and nonlinear dependencies. Similarly, if we remove the liner effect, the correlation will go to zero whereas the mutual information will capture the true residual nonlinear dependencies between two distributions.

### 3.2. On fMRI Data

We measured the nonlinear dependencies among 47 component time courses estimated from the resting fMRI data. The results are shown in **Fig. 2**. Panel A is the average FNC over 314 subjects of 47 components. The average of nonlinear dependencies of 47 components is calculated via our proposed MI method for each of the 314 subjects. The result of applying T-test for one sample to show the difference of the average value from minimum average is presented in **Fig. 2**. B. The FDR threshold (p < 0.05) shows that most of the differences relative to the minimum mean for all entries in this matrix are significant. This shows us the degree to which MI differences are significantly different across the FNC matrix and demonstrates modular nonlinear dependencies between visual (VIS) and somatomotor (SM) components with other components. Interestingly, despite high linear correlation between subcortical (SC) and auditory (AUD), a low rate of nonlinear dependency is observed among them.

**Fig. 2.**
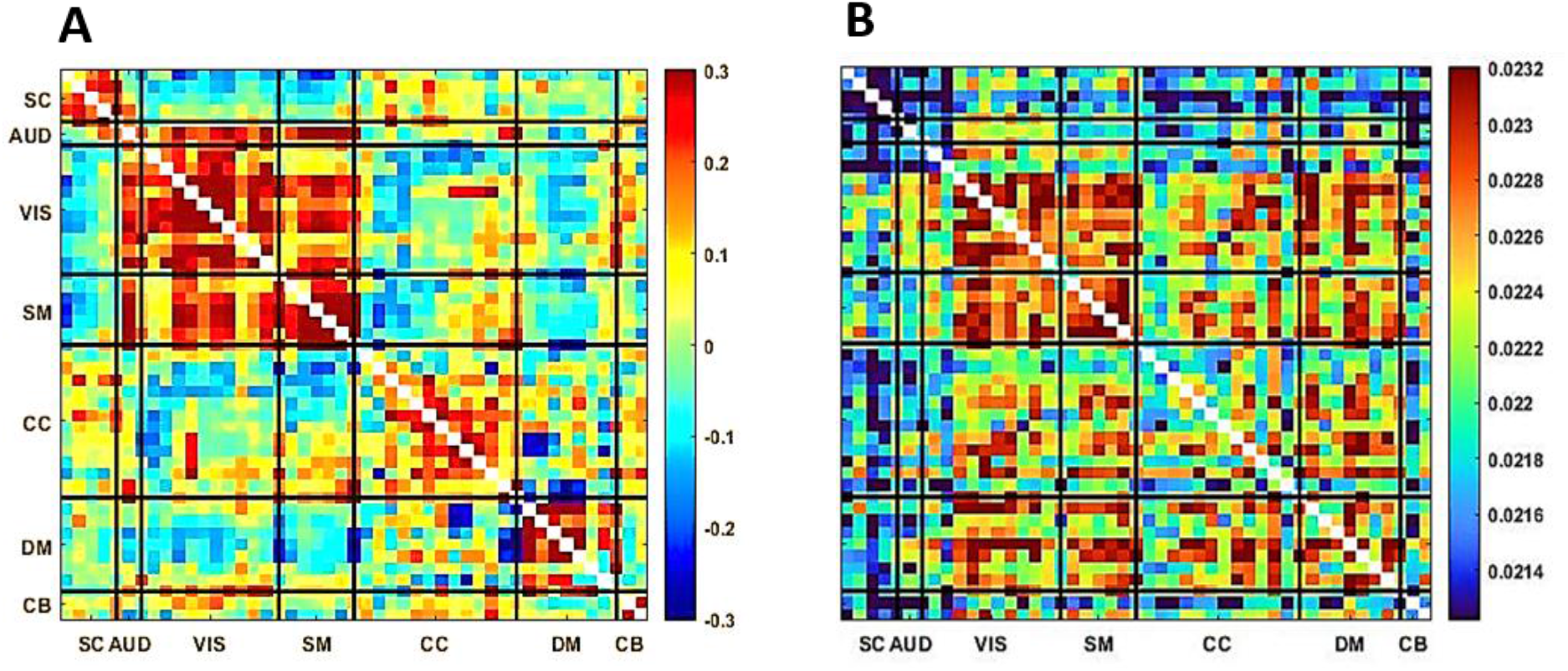
**A)** Mean (linear) functional network connectivity (FNC) over 314 subjects. **B)** Mean mutual information (MI) after removing the linear correlation over 314 subjects. Note that both linear and nonlinear effects are modularized, but in different ways suggesting they are providing complementary information.

Next the variation in nonlinear dependencies between healthy controls and schizophrenia patients are evaluated using our MI method and compared using a two-sample T-test. In **Fig. 3**, panel A, the lower triangle shows −*log*10(*p*) × *sign*(*T*) before threshold multiply by the initial p-value before FDR. The upper triangle is the same as lower one, but p-values are threshold such that entries shown in a different color represent significant differences in nonlinear dependency between groups. We observe a significant difference in nonlinear dependencies between visual (VIS) components to other components such as auditory (AUD), visual (VIS), somatomotor (SM), cognitive control (CC), and default-mode (DM) in schizophrenia (SZ) patients relative to healthy controls (HC). We also visualized differences in the joint density among the most nonlinear networks and identified several interesting patterns that would be completely missed in a typical linear analysis. Panel B, the connectogram shows components with significant difference in their nonlinear dependencies between healthy control and schizophrenia patients such that two components are connected with a line if the difference in their nonlinear dependencies between HC and SZ is significant (with yellow for HC>SZ and blue for SZ>HC).

**Fig. 3.**
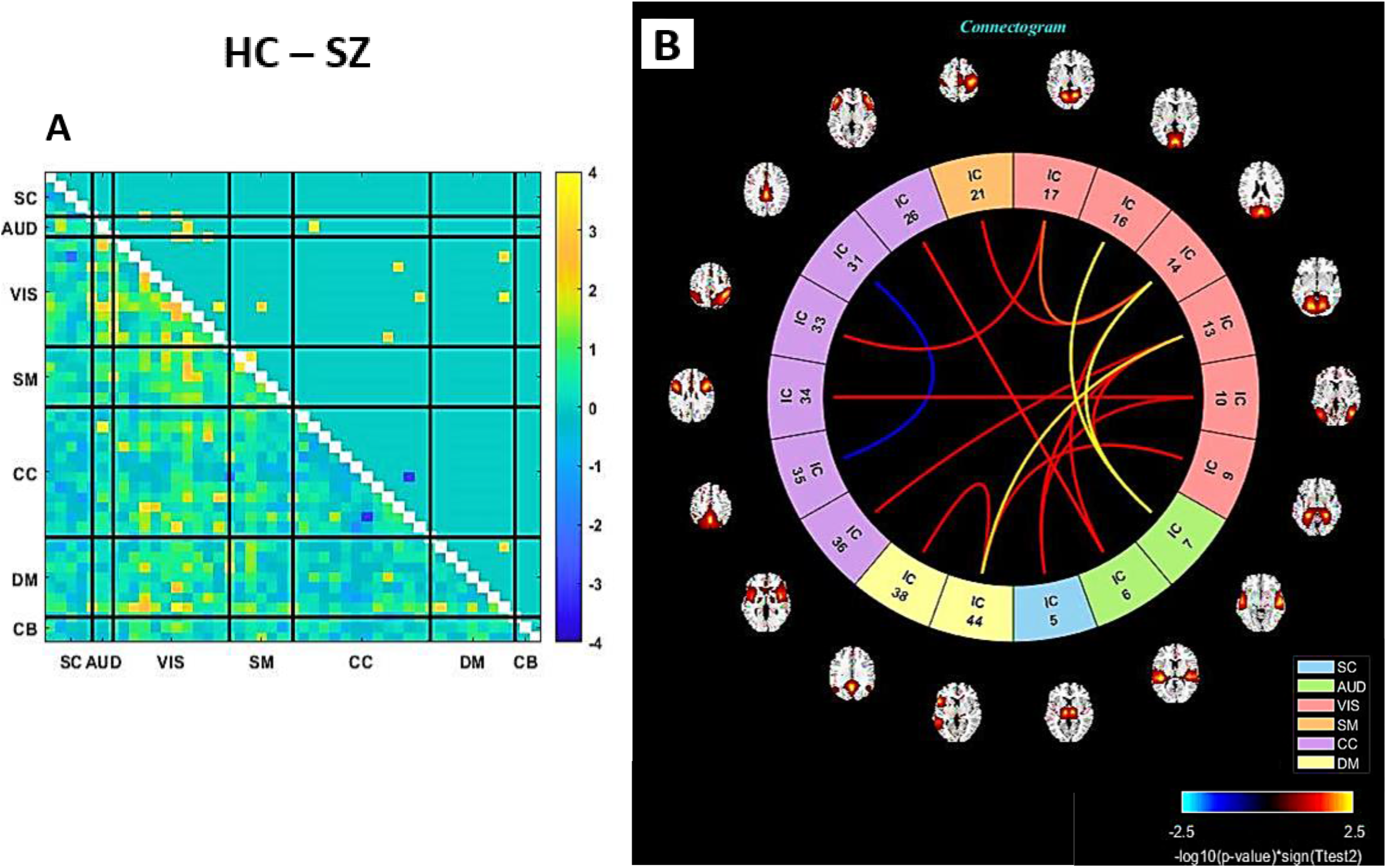
**A)** Upper triangle: The group difference (HC-SZ) in MI after removing the linear correlation. The p-values are adjusted by FDR and threshold (p < 0.05). Values are plotted as −log10(p-value) × sign(t-statistics). Lower triangle: Identical to the upper tringle except that p-values are not threshold, also the values are multiplied by the initial p-value before FDR. **B)** Connectogram that show components with significant nonlinear correlation in HC- SZ are connected. In all but one case, results show significantly more nonlinearity in the controls, mostly linked to the visual domain.

### 3.3. Boosted Approach

Dependencies among components in healthy controls and schizophrenia patients are assessed with three methods: 1. Pearson correlation, in which the emphasis is on only linear correlation, 2. Mutual information as described in 2.3. *A Mutual Information Approach*, is quantifying only nonlinear dependencies, and 3. the boosted approach explained in *2.6. Boosted Approach*is boosting the linear correlation by capturing nonlinear dependencies.

Next, in each method, T-test is applied to compare the differences between two groups. Adjusted p-value by FDR is threshold (p<0.05). The number of pairs with significant differences for Pearson correlation method is 530, in mutual information method is 17 and in Boosted method is 537.

### 3.4. Joint Distributions

We were also interested in visualizing the relationship among the timecourse pairs which exhibited nonlinear differences between patients and controls. **Fig. 4** demonstrates the differences between healthy controls and schizophrenia patients for the five pairs showing the largest group differences. Values increases from left to right and up to down. Panel A shows the difference of joint distribution of the 26^th^ and 6^th^ components. The 26^th^ component belongs to cognitive control (CC) 6^th^ component is in the auditory (AUD) domain. Panel B is the joint distribution difference of the 44^th^ and 13^th^ components. The 44^th^ component is in default-mode (DM) and 13^th^ component is in visual (VIS) domain. Panel C represent the joint distribution difference of 7^th^ and 16^th^ components. The 7^th^ component is in auditory (AUD) and 16^th^ component is in visual (VIS) domain. Panel D illustrates the difference of joint distribution of 14^th^ and 7^th^ components. Panel E exhibits the joint distribution of 14^th^ and 17^th^ components. Both 14^th^ and 17^th^ components belong to visual (VIS) domain. Panels B, D, and E share some similarities, including a negative relationship between the two components.

**Fig. 4.**
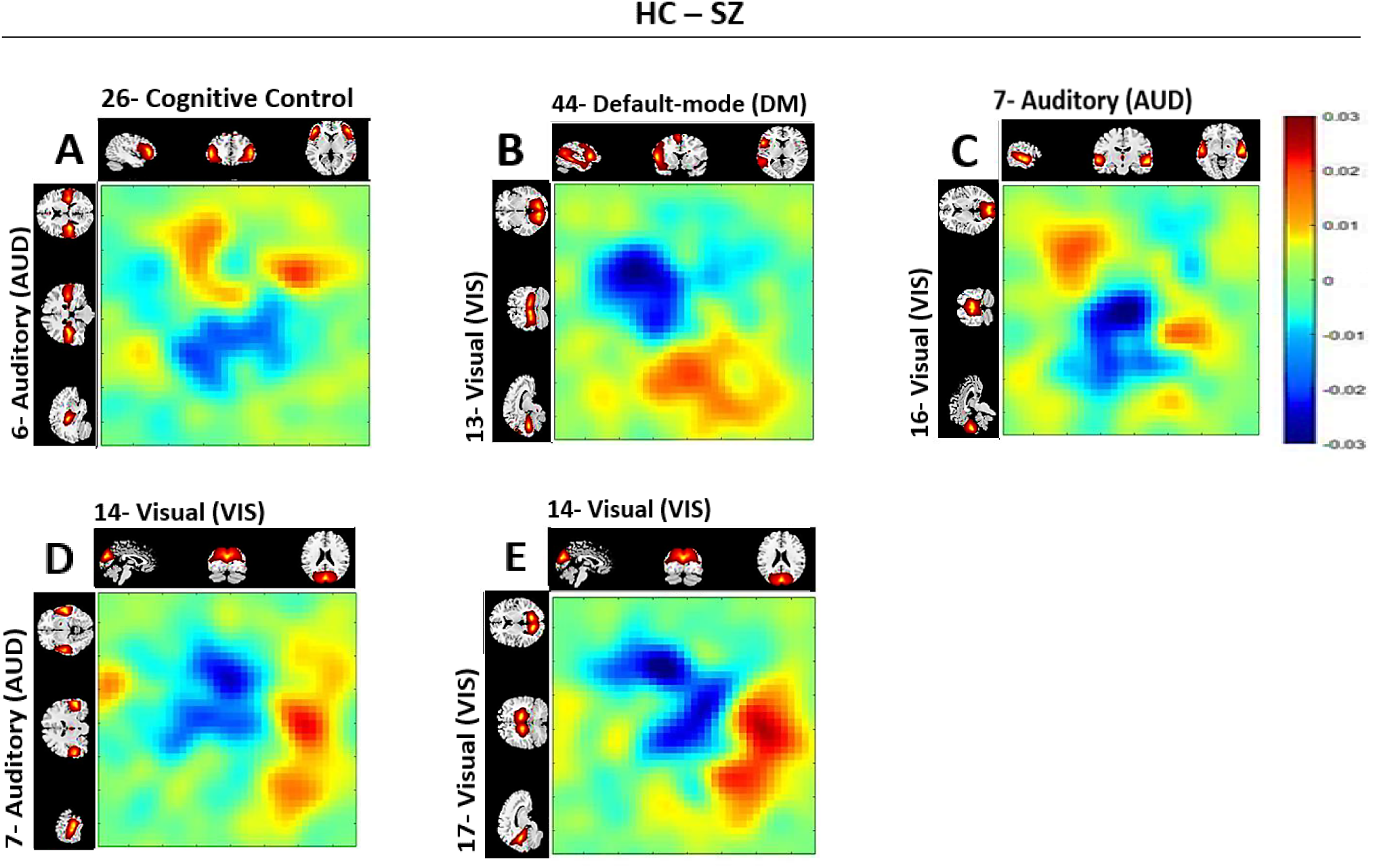
Difference between the HC and SZ joint distributions for the five pairs showing the largest group differences in the nonlinear dependencies of pairs (26, 6), (44, 13), (7, 16), (14, 7), (14, 17). Values in each distribution increases from left to right and up to down. The 26th component belongs to cognitive control (CC). The 6th and 7th components are in auditory (AUD). The 44^th^ component is in default-mode (DM). All 13^th^, 14^th^, 16^th,^ and 17^th^ components are in visual (VIS) domain. We observe some interesting differences in the joint distributions with patients generally showing differentially higher activity in one network and lower activity in the paired network.

In **Fig. 4**. A, we can observe controls spend more time in low level of an auditory component #6 relative to SZ regardless of the values of the 26^th^ components. From B, healthy controls are considerably more active in both the default model network (#44) and a visual component (#13) than in SZ. In panel C, we notice healthy controls show less activity in both an auditory (#7) and visual (#16) components compare to SZ. From D and E, it can be interpreted those healthy controls show a higher level of activation in visual (#14) components relative to a more posterior visual (#17) and an auditory (#7) component than do the patients.

For these 5 pairs, nonlinear FNC shows the hyper-connectivity compared to SZ i.e., shows the nonlinear part of two distributions has more dependency in HC. However, the plot of group differences in the joint distribution can illustrate how two pairs that show high dependency in HC, can be differently distributed in SZ. For example, in **Fig. 4**, we can observe how differently these pairs in SZ and HC are distributed. Panel B, D, and E, healthy control time course have higher levels of activation (red color is dragged to the lower corner and blue to the top left) while in panels A and C, health controls have lower activation level (red is dragged to the upper left and blue to the lower right).

## Discussion

In this preliminary work we highlight the benefit of studying nonlinearities in functional connectivity. Previous functional connectivity studies are based on correlation coefficient that assess the linear correlation only, and as a result they miss the nonlinear contributions. We establish an approach to assess the explicitly nonlinear dependencies between distinct regions of the brain by first removing the linear dependencies. We first demonstrated our approach works as expected on simulated data (**Fig. 1**). Following the nonlinear dependencies among 47 timecourses on 314 subjects are assessed (**Fig. 2**). A similar approach was applied to estimate how differently in average distinct regions of a schizophrenia patient’s brain contributes nonlinearly to the context of functional connectivity (**Fig. 3**). Also, the joint distribution of five pairs with the largest group differences in the nonlinear dependencies in HC-SZ is studied (**Fig. 4**).

There are a number of possible causes of nonlinear dependencies including: 1) Nonlinear hemodynamic effects. Studies on the relationship between neuronal activity, oxygen metabolism, and hemodynamic responses have shown the link between neuronal activity and hemodynamic response magnitude exhibits both linear and nonlinear effects in task data (cite fMRI nonlinear task study). Other results suggest a strongly nonlinear relationship between electrophysiological measures of neuronal activity and the hemodynamic response (Devor et al., 2003; Sheth et al., 2004), 2) Differences in blood flow, blood oxygenation, and blood volume both within subjects and between groups. Experiments indicates that acquires vascular space occupancy (VASO), arterial spin labeling (ASL) perfusion and BOLD signals respond nonlinearly to stimulus duration (Gu, Stein, & Yang, 2005) 3) Subject motion. Even minor head posture changes may result in considerable spatially complex field changes in the brain (Liu, de Zwart, van Gelderen, Murphy-Boesch, & Duyn, 2018). While we cannot completely exclude motion, we carefully curated the data to focus on low motion subjects and in addition there were no significant motion differences between the groups(Damaraju et al., 2014).

The different modularized patterns evident in linear and nonlinearly modularity suggests a complementarity of the nonlinear and linear relationships. It may be important to capitalize on these differences in future studies. Our results suggest interesting variation among networks. For example, as shown in **Fig. 2**. B, significant nonlinear dependencies are observed between visual (VIS), somatomotor (SM) domains and within cognitive control (CC) and default-mode (DM) domains. The auditory (AUD) network shows strong differences in linear dependencies (A), but not much nonlinear, whereas both visual and sensorimotor show strong within domain nonlinear dependencies (B). Also, relatively low rate of nonlinear dependencies is observed between subcortical (SC) and auditory (AUD) with other components.

We also found significant differences in the nonlinear relationships among the patients and controls. Nonlinear FNC pairwise comparison between SZ and HC are shown in **Fig. 3**. Part A. In most cases the controls are showing higher nonlinear dependencies relative to patients, mostly linked to the visual domain. There is a significant difference in nonlinear correlation within visual (VIS) components as well as between VIS components and to other components such as auditory (AUD), somatomotor (SM), cognitive control (CC), and default-mode (DM) in SZ patient and HC. We observe that most of the patient/control differences involve visual and auditory components. This is intriguing given existing evidence suggesting some schizophrenia symptoms may be linked to the visual system (Gong et al., 2020; Johnston, Stojanov, Devir, & Schall, 2005; Onitsuka et al., 2007). Having said that, visual symptoms such as visual hallucinations are rather uncommon in SZ, and rarer than auditory and tactile abnormalities (van de Ven, Rotarska Jagiela, Oertel-Knöchel, & Linden, 2017). In addition, some studies suggest inborn blindness may be shielding against the development of schizophrenia, characterized by inevitably noisy perceptual input that causes false inferences. These finding argue that when individuals cannot see from birth, they depend more on the other senses. Thus, the resulting model of the world is more resistant to false interpretations (Landgraf & Osterheider, 2013; Leivada & Boeckx, 2014; Morgan et al., 2018; Pollak & Corlett, 2019; Riscalla, 1980).

There is still much work to be done. Future work should focus on carefully evaluating the possible sources of the nonlinear relationships. Quantitative fMRI studies could be used to isolate nonlinearities in blood oxygenation, volume, and flow. In addition, high field layer specific fMRI studies could be used to evaluate nonlinearities in input vs output layers. The contribution of various physiological variables (e.g., respiration, CO_2_, heart rate, and motion) could also be evaluated in future work.

In sum, our results provide evidence suggesting there are meaningful and significant nonlinear dependencies among fMRI time courses. We have showed evidence suggesting there are meaningful (modularized and group different) super-linear effects in FNC which primarily implicates the visual cortex as disrupted in schizophrenia. We present two approaches, a focus on the explicitly linear effects or a boosted approach which captures both linear and nonlinear effects within one metric. Future work should further study information contained in the nonlinear relationships, and could be studies with faster acquisitions, linked to multimodal imaging such as concurrent EEG data, and replicate the results we show in this work.

## A. Appendix

## Supplementary Data

Supplementary data to this article can be found online at http://dx.doi.org/10.1016/j.nicl.2014.07.003

